# Myc upregulates Ggct, γ-glutamylcyclotransferase to promote development of *p53*-deficient osteosarcoma

**DOI:** 10.1101/2024.03.18.585455

**Authors:** Tomoya Ueno, Shohei Otani, Yuki Date, Yu Katsuma, Yuma Nagayoshi, Tomoko Ito, Hiromi Ii, Susumu Kageyama, Susumu Nakata, Kosei Ito

## Abstract

Osteosarcoma (OS) in humans is characterized by alterations in the *TP53* gene. In mice, loss of p53 triggers OS development, for which c-Myc (Myc) oncogenicity is indispensable. However, little is known about which genes are targeted by Myc to promote tumorigenesis. Here, we examined the role of Ggct, γ-glutamylcyclotransferase which is a component enzyme of γ-glutamyl cycle essential for glutathione homeostasis, in human and mouse OS development. We found that *GGCT* is a poor prognostic factor for human OS, and that deletion of *Ggct* suppresses *p53*-deficient osteosarcomagenesis in mice. Myc upregulates Ggct directly by binding to the *Ggct* promoter, and deletion of a Myc binding site therein by genome editing attenuated the tumorigenic potential of *p53*- deficient OS cells. Taken together, these results show a rationale that GGCT is widely upregulated in cancer cells and solidify its suitability as a target for anticancer drugs.

## INTRODUCTION

Osteosarcoma (OS) is a malignant bone tumor for which few targeted therapies are effective. In humans, the frequent inactivation of *TP53* in sporadic OS^1,2^, as well as its germline mutation in cases of Li-Fraumeni syndrome with a high incidence of OS^3,4^, provide robust evidence for the role of p53 as a critical tumor suppressor of OS development. In mice, loss of p53 in osteoprogenitor or mesenchymal stromal cells (MSCs) is sufficient to trigger osteosarcomagenesis; this was well-documented using the *Osterix (Osx)*/*Sp7*-Cre; *p53*^fl/fl^ mouse model ^5,6^. In a series of previous studies^7–9^, we identified a vital oncogenic axis comprising Runx3, a member of Runx family of transcription factors, and c-Myc (Myc), a pivotal tumor-promoting factor in OS^10^. In the absence of p53, Runx3 aberrantly upregulates Myc in human and mouse OS cells^7^. A better understanding of the precise molecular mechanism underlying pathogenesis of *p53*- deficient OS will be achieved by identifying the genes targeted by the Myc transcription factor; however, the genes that are both targeted by Myc and are essential for development of OS remain unknown

Glutathione (GSH) is γ-glutamyl-cysteinylglycine, which is biosynthesized from glutamate, cysteine, and glycine^11^. It is the most abundant antioxidant *in vivo* and has multiple functions; however, in cancers, excessive elevation of GSH promotes tumor progression^12^. GGCT, γ-glutamylcyclotransferase, originally named as C7orf24, is an enzyme involved in the γ-glutamyl cycle that is essential for GSH homeostasis, and is known for its oncogenicity and utility as a tumor marker^13,14^. It is thought that the first detection of elevated GGCT in tumors was a study comparing bladder urothelial carcinomas and normal controls^15^, followed by reports of elevated levels in breast, ovarian, cervical, lung, colon, and esophageal squamous cell carcinomas, glioma, and OS^16^. In human OS, the levels of GGCT were considerably higher both in cell lines and primary tumors and *GGCT*-knockdown reduces tumor cell growth, invasiveness and motility^17^. The cell cycle-dependency of *GGCT* promoter activation suggests that upregulation of GGCT plays a role in cancer cell proliferation^18^. In fact, a recent study shows that Ggct is upregulated downstream of Kras oncogenic signaling in a mouse model of lung cancer, indicating its pivotal role in the production and metabolism of GSH, which is critical for promotion of carcinogenesis^19^. However, we do not know what is driving Ggct expression as the transcription factors remain unknown.

Here, we show that Myc, the most widely functional oncogenic transcription factor in human cancers^20^, targets and directly upregulates Ggct. GGCT is upregulated significantly in patients with OS and is a poor prognostic factor. In addition, loss of Ggct suppresses the increase in GSH levels associated with tumorigenesis, and attenuates development of OS in *p53*-deficient mice. Finally, deletion of the Myc-binding site from the *Ggct* promoter region by genome editing reduced tumorigenicity of *p53*-deficient OS cells. Taken together, these results demonstrate that GGCT is an attractive target for development of new anticancer drugs, which is what we are aiming to do^21,22^.

## RESULTS

### GGCT is upregulated in human OS and is a poor prognostic factor

Biosynthesis of GSH requires sufficient amounts of glutamate, cysteine, and glycine to maintain appropriate levels of the tripeptide. The γ-glutamyl cycle is thought to play a central role in GSH homeostasis by transporting amino acids such as glutamate, cysteine, and glycine^13^ (Figure 1A). This cycle involves enzymes; two ATP-dependent ligases, glutamate cysteine ligase (GCL) and glutathione synthase (GSS), both of which are essential for GSH synthesis, γ-glutamyltranspeptidase (GGT), 5-oxoprolinase (OPLAH), and GGCT (Figure 1A).

**Figure 1.**
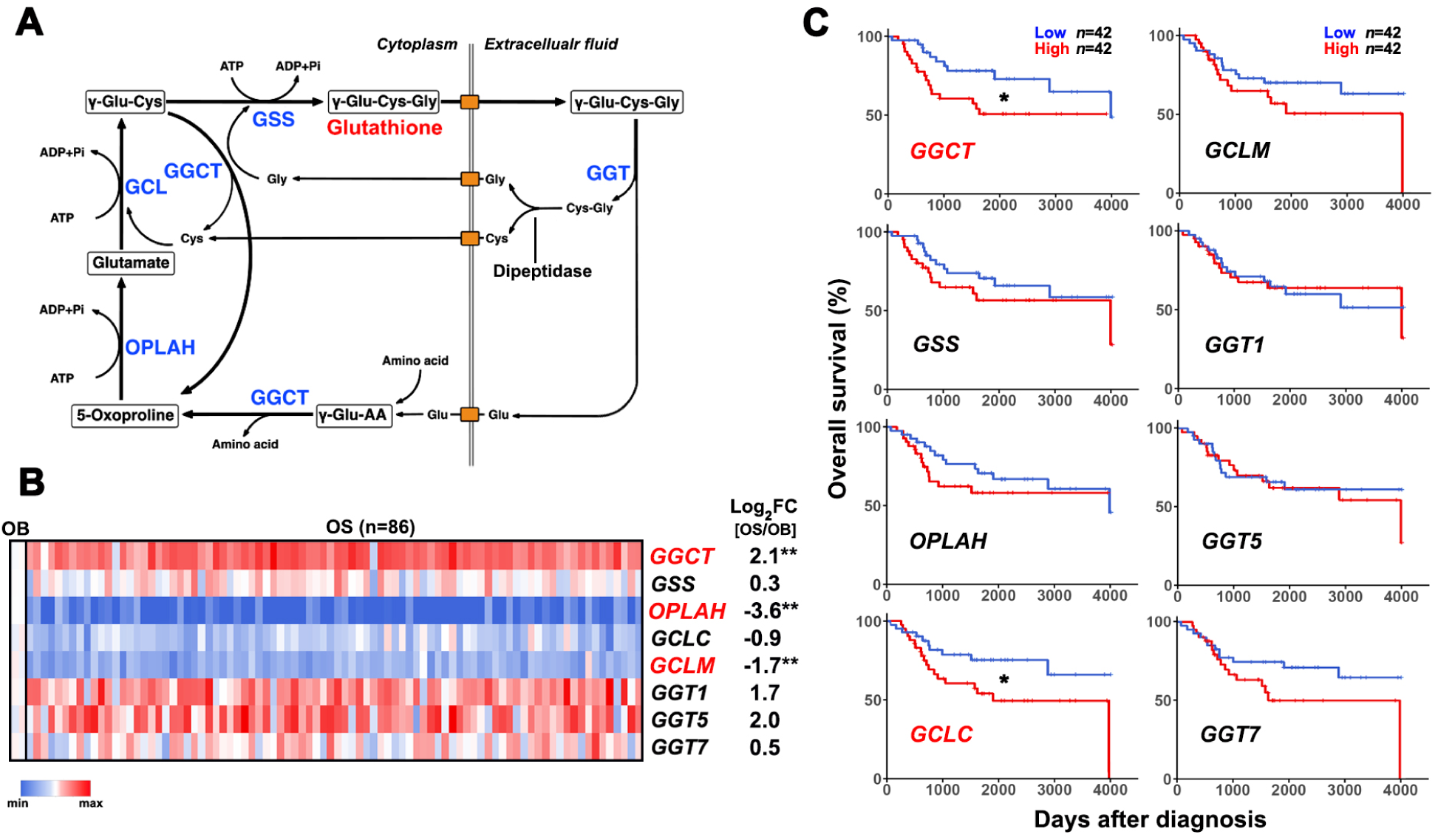
Upregulation of GGCT in human OS is an unfavorable prognostic factor. (A) Overview of the γ-glutamyl cycle and its component enzymes. GGCT, γ- glutamylcyclotransferase; GGT, γ-glutamyltranspeptidase; GCL, glutamate cysteine ligase; GSS, glutathione synthetase; OPLAH, 5-oxoprolinase; γ-Glu-Cys-Gly; glutathione. (B) Heatmap showing color-coded gene expression levels of γ-glutamyl cycle enzymes across 86 OS patients. The ratio of gene expression level in OS versus normal osteoblast cells (OB) is shown as log_2_FC. GCL comprises a catalytic subunit (GCLC) and a modifier subunit (GCLM). \*\**p*<0.01. (C) Prognostic value of γ-glutamyl cycle enzyme genes, as determined by Kaplan–Meier survival analysis of OS patients from the TARGET cohort. \**p*<0.05.

To examine the involvement of these enzymes in development of human OS, we first compared the transcriptome of human OS tissues with that of normal human osteoblasts. The Therapeutically Applicable Research to Generate Effective Treatments (TARGET) cohort was used to evaluate changes in expression of genes encoding the enzymes. Inactivation of p53 is critical for development and progression of OS in both humans and mice^8^. Almost all cases (84 of 86) in the TARGET cohort harbored gene alterations of *TP53*^7^. Among genes encoding the component enzymes, *GGCT* was significantly upregulated, and *OPLAH* and *GCLM* were downregulated, in OS tissues (Figure 1B). Of these, only *GGCT* was associated with a poor prognosis in the human OS cohort (Figure 1C). *GCLC* was also a significant poor prognostic factor (Figure 1C), although its expression fell in OS patients (Figure 1B). These results suggest that GGCT within the γ-glutamyl cycle is a tumor-promoting factor for human OS.

### Ggct is upregulated in *p53*-deficient mouse OS cells

Loss of p53 in osteoprogenitor or MSCs is sufficient for OS development, indeed, *Osx*/*Sp7*-Cre; *p53*^fl/fl^ mice (hereafter referred to as *OS* mice) are used widely to study the molecular mechanism underlying OS development^5–7^. When we compared expression of genes encoding the eight enzymes in cells comprising OS tissue in *OS* mice (i.e., mOS-1, 2, and 3), we found that Ggct was elevated significantly in mOS cells compared with MSCs (i.e., MSC-1, 2, and 3) used as controls (Figure 2A). Higher amounts of Ggct protein was also immunodetected in mOS cells than in MSCs (Figure 2B); accordingly, the amount of GSH in mOS cells increased (Figure 2C). The tumorigenic potential of clonal mOS cells (derived from mOS-1 and mOS-2 cells) transplanted into immunodeficient mice (i.e., allograft) was proportional to the expression level of Ggct (Figure 2D and 2E). These results suggest that Ggct functions as an oncogene in *p53*- deficient OS in mice in the same way as in human OS.

**Figure 2.**
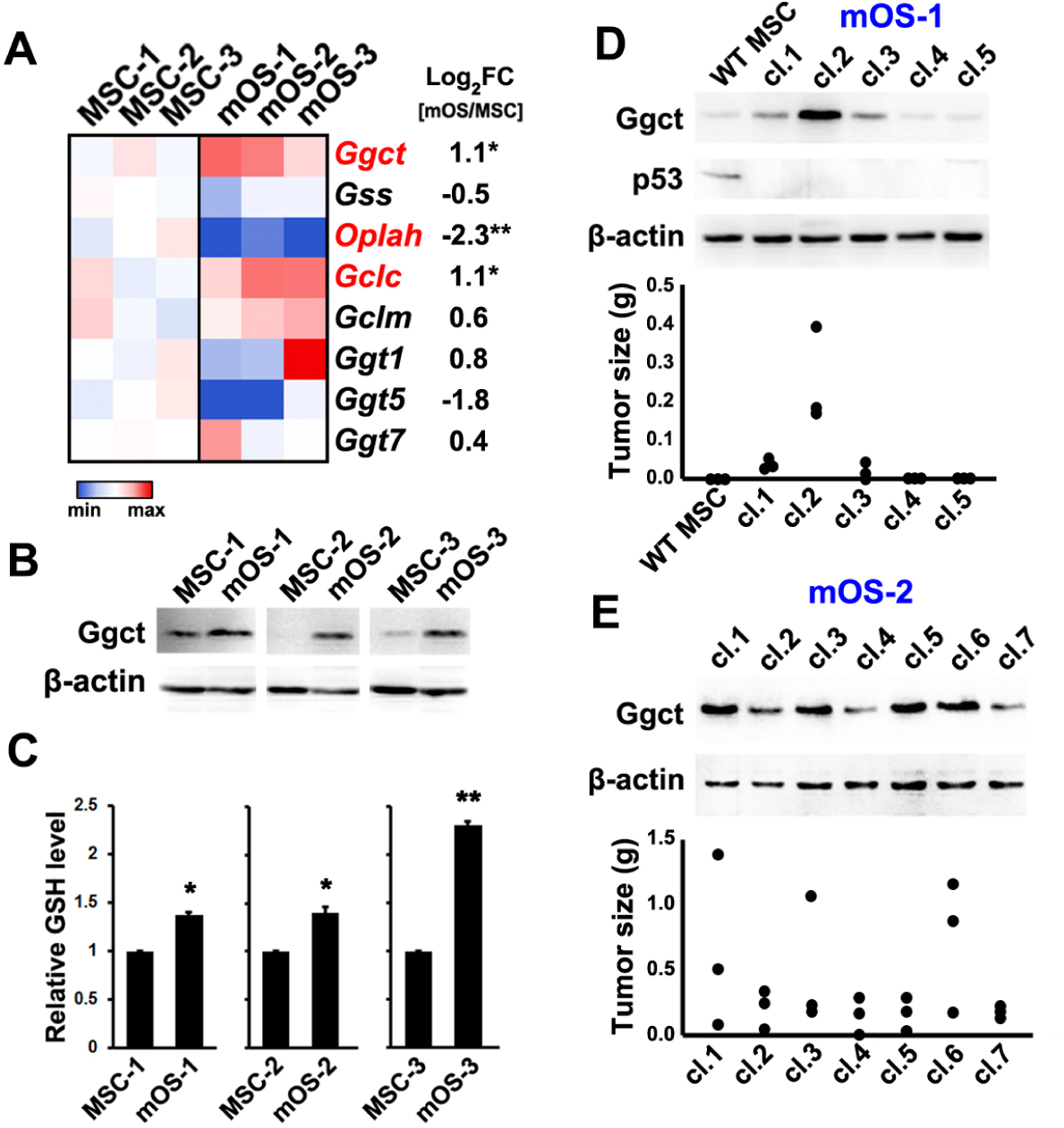
Ggct is upregulated in *p53*-deficient mouse OS cells. (A) Heatmap showing color-coded expression levels of genes encoding γ-glutamyl cycle enzymes in three sets of MSCs (MSC-1, 2, 3) and mOS cells (mOS-1, 2, 3) isolated from three individual *OS* mice. The ratio of the gene expression level in mOS cells to that in MSCs is shown as the log_2_FC. (B) Western blot analysis of the indicated proteins in MSCs and mOS cells. (C) Relative amounts of GSH in MSCs and mOS cells. Data are presented as the mean ±SE (*n*=3). \*\**p*<0.01; \**p*<0.05. (D and E) Western blot analysis of the indicated proteins in clonal mOS cells isolated from OS derived from two individual *OS* mice, mOS-1 (D) and mOS-2 (E), and MSCs from a 1-year-old wild-type mouse (WT MSC) in D. The tumorigenicity of each clone was evaluated by allograft using nude mice (n = 3) (D and E).

### Ggct is oncogenic in *p53*-deficient OS

To evaluate the oncogenic role of *Ggct* during development of *p53*-deficient OS *in vivo*, we generated a *Ggct*-disrupted mouse line based on homologous recombination in ES cells (Figure S1). Deletion of *Ggct* prolonged the life span of *OS* mice and reduced the incidence of OS (*OS* mice vs *OS*; *Ggct*^-/-^ mice) (Figure 3A and 3B). In addition, compared with MSCs from *OS* mice, there was an increase in GSH levels in OS cells (Figure 2C), but not in *OS*; *Ggct*^-/-^ mice (Figure 3C). These observations strongly suggest that Ggct is a tumor-promoting factor in *p53*-deficient OS cells and is responsible for increased expression of GSH during tumorigenesis *in vivo*.

**Figure 3.**
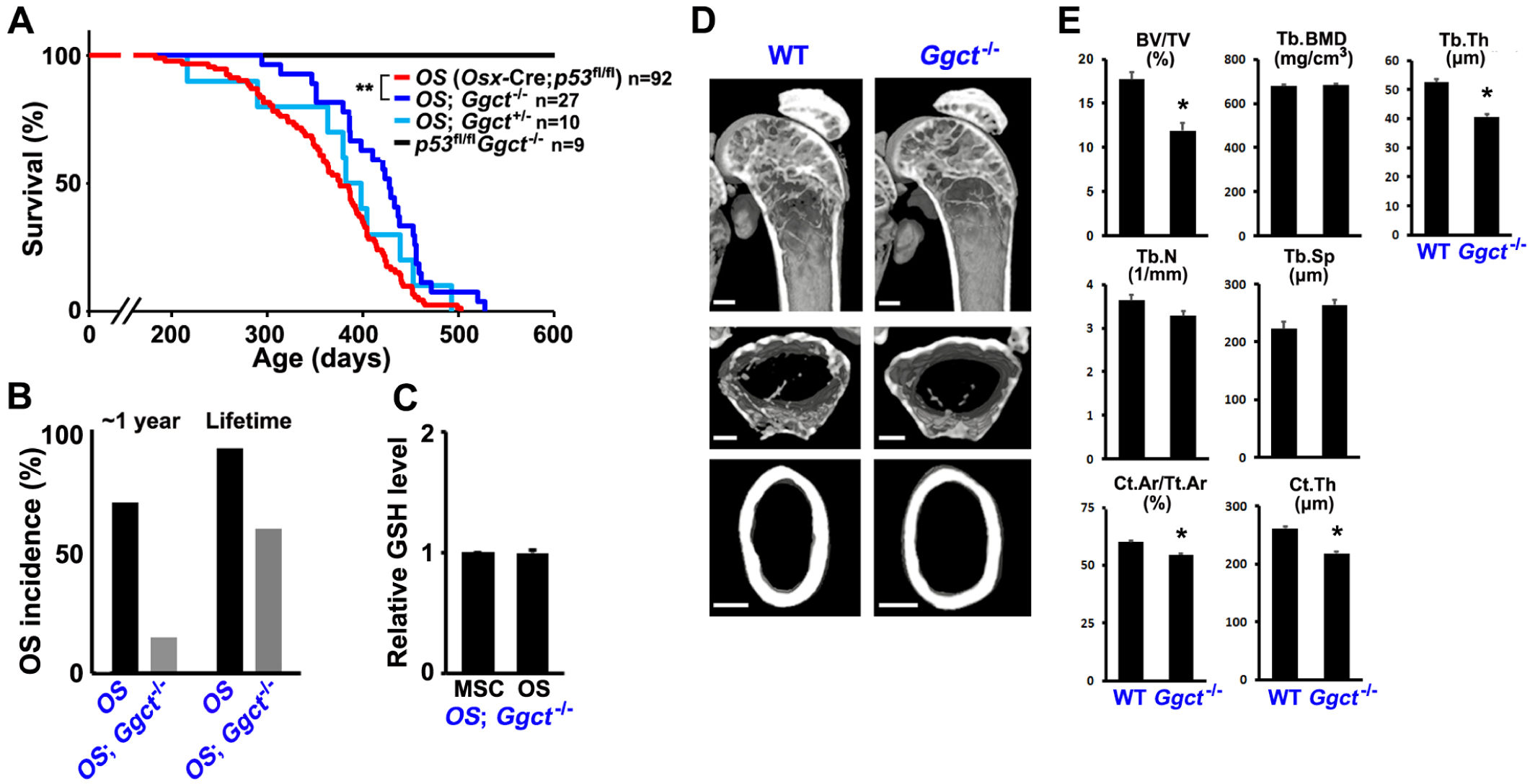
Ggct is oncogenic in *p53*-deficient OS. (A) Survival of *Ggct*^+/+^, *Ggct*^+/-^, and *Ggct*^-/-^ mice on the *OS* (*Osx*-Cre; *p53*^fl/fl^) background, alongside Cre-free controls. \*\**p*<0.01. (B) Incidence of OS in the indicated genotypes at 1 year-of-age and throughout life. (C) Relative amounts of GSH in MSCs and mOS cells of *Ggct*^-/-^ mice on the *OS* background (*OS*; *Ggct*^-/-^ mice). Data are presented as the mean ±SE (*n*=3). (D) Representative three- or two-dimensional µCT images of the bone architecture of male wild-type (WT) and *Ggct*^-/-^ mice at 6 months-of-age. Images of trabecular bone at the distal femoral metaphysis (upper and middle) and cortical bone at the mid-diaphysis in femurs (lower). Scale bars = 500 µm. (E) Quantification of the trabecular bone volume (bone volume/tissue volume, BV/TV), trabecular bone mineral density (Tb.BMD), trabecular thickness (Tb.Th), trabecular number (Tb.N), trabecular separation (Tb.Sp), cortical area (Ct.Ar/Tt.Ar), and cortical thickness (Ct.Th) in male WT and *Ggct*^-/-^ mice aged 6 months. Data are presented as the mean ±SE (*n*=3). \**p*<0.05.

Systemic deletion of *Ggc*t did not affect the life span of mice (*p53*^fl/fl^*Ggct*^-/-^ mice in Figure 3A), in line with observations reported previously^19^. However, micro-CT (µCT) analysis revealed that deletion of Ggct reduced trabecular and cortical bone formation in the femur (a common site of OS) of male and female adult mice, although the trend was mild and was not observed for all parameters examined (Figures 3D, 3E, S2A and S2B). Therefore, Ggct may play a physiological role in the promotion of bone formation.

### Myc upregulates Ggct in *p53*-deficient OS

The data obtained thus far show a significant positive correlation between Ggct expression and Myc expression in OS cells; mOS cell clones with high Ggct expression and tumorigenicity (Figures 2D and E) also show higher levels of Myc expression (Figures 4A and B). Knockdown of Myc in mOS cells reduced expression of Ggct (Figure 4C). MYC knockdown in human OS cell lines MNNG-HOS, NOS1, and Saos-2 also reduced expression of GGCT (Figure 4D); furthermore, analysis of human OS patients revealed that among all genes encoding enzymes in the γ-glutamyl cycle, *GGCT* showed the strongest correlation with *MYC*, with a Spearman’s rank correlation coefficient (R) of 0.46 (Figure 4E and Figure S3). This was also supported by immunohistochemical analyses, which showed that cells with a Myc-positive nucleus also had Ggct-positive cytoplasm in OS tissues from two individual *OS* mice, mOS-1 and mOS-2, from which mOS-1 and mOS-2 cells shown in Figure 2 were derived, respectively (Figure 4F). Taken together, these results suggest that MYC/Myc upregulates GGCT/Ggct in both human and mouse OS cells.

**Figure 4.**
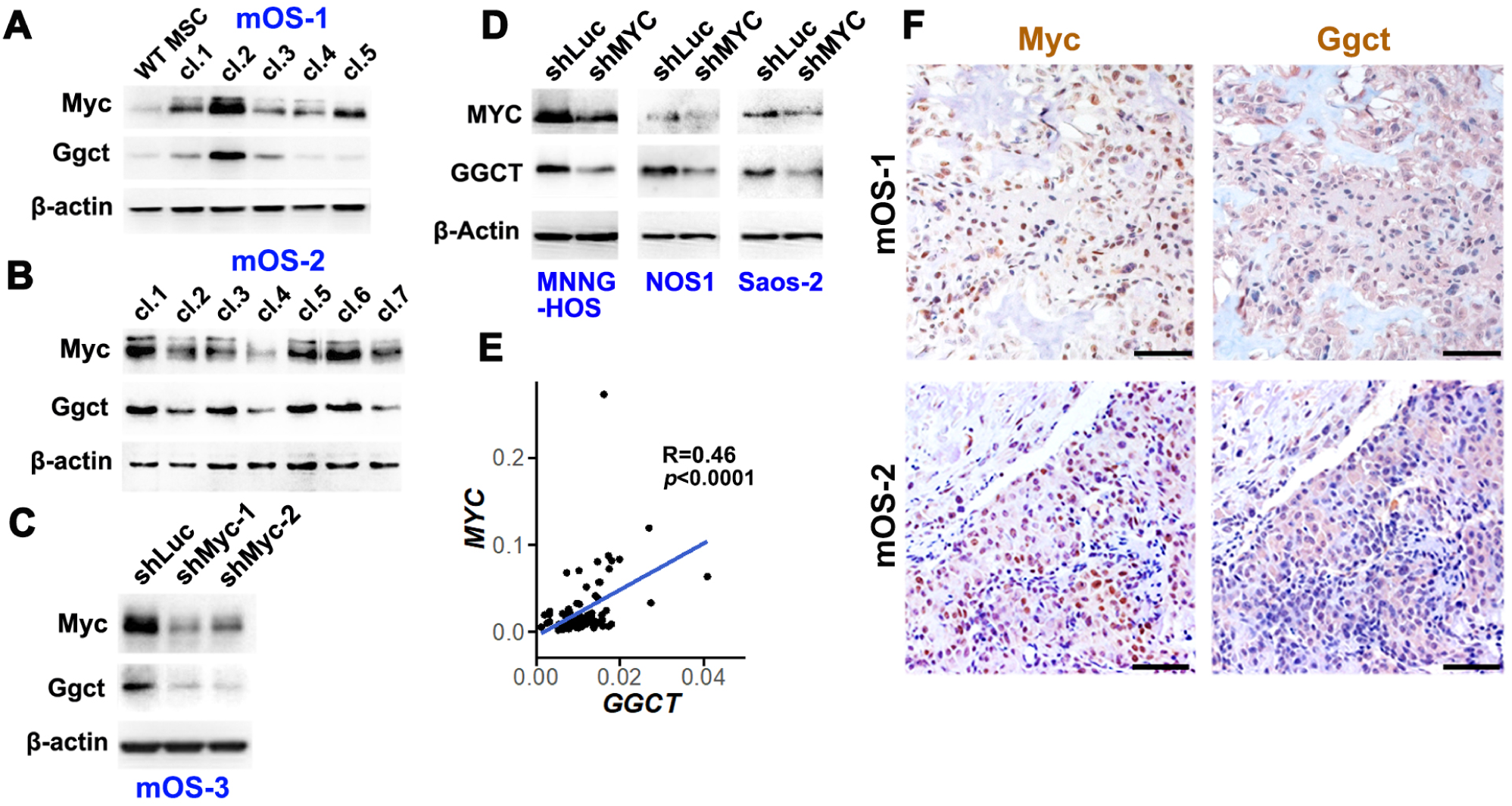
Myc upregulates Ggct in *p53*-deficient OS. (A and B) Western blot analysis of the indicated proteins in clonal mOS cells isolated from OS formed in two individual *OS* mice, mOS-1 (A) and mOS-2 (B), and MSCs from a 1-year-old wild-type mouse (WT MSC) (A). The immunoblots showing expression of Ggct and β-actin in A and B, respectively, are identical to those shown in Figure 2D and 2E, respectively. (C) Effect of knockdown of Myc on expression of Ggct in mOS-3 cells, as determined by western blotting. (D) Effect of MYC knockdown on expression of GGCT in human OS cell lines; MNNG-HOS, NOS1 and Saos-2, as determined by western blotting. (E) MYC expression levels plotted against GGCT expression levels in human OS (n=86, Figure 1B). Spearman’s rank correlation coefficient (R) is shown. (F) Immunodetection of Myc and Ggct in mOS-1 and mOS-2 cells. Left and right panels show serial sections. Counterstaining was done with hematoxylin. Scale bars = 50 µm.

### Myc-mediated upregulation of Ggct is critical for tumorigenicity of *p53*-deficient OS cells

To examine the mechanism by which Myc regulates Ggct in more detail, we investigated the genome-wide profiles of Myc occupancy and chromatin activation using ChIP-seq analysis of Myc and H3K4me3, respectively, as well as an analysis of open chromatin using the assay for transposase-accessible chromatin using the sequencing (ATAC-seq) method. In the *Ggct*-coding region in the mouse genome, only its promoter was found to be chromatin-activated, open, and occupied by Myc (Figure S4). The *Ggct* promoter harbors a consensus binding site for Myc. (Figure 5A). In human OS cells, similarly, MYC occupied the *GGCT* promoter and its chromatin was activated and open (Figure S5A), however, we observed no significant occupancy of the *GGCT*-coding region by MYC; the only exception was the promoter (data not shown). Occupancy of the *GGCT* promoter by MYC was comparable with occupancy of the *CDK4* promoter, another target gene for MYC^23^, in Saos-2 cells (Figure S5B), in which GGCT was downregulated by MYC knockdown (Figure 4D). Therefore, we focused on the role of Myc at the Myc-consensus site in the *Ggct* promoter of mOS cells.

**Figure 5.**
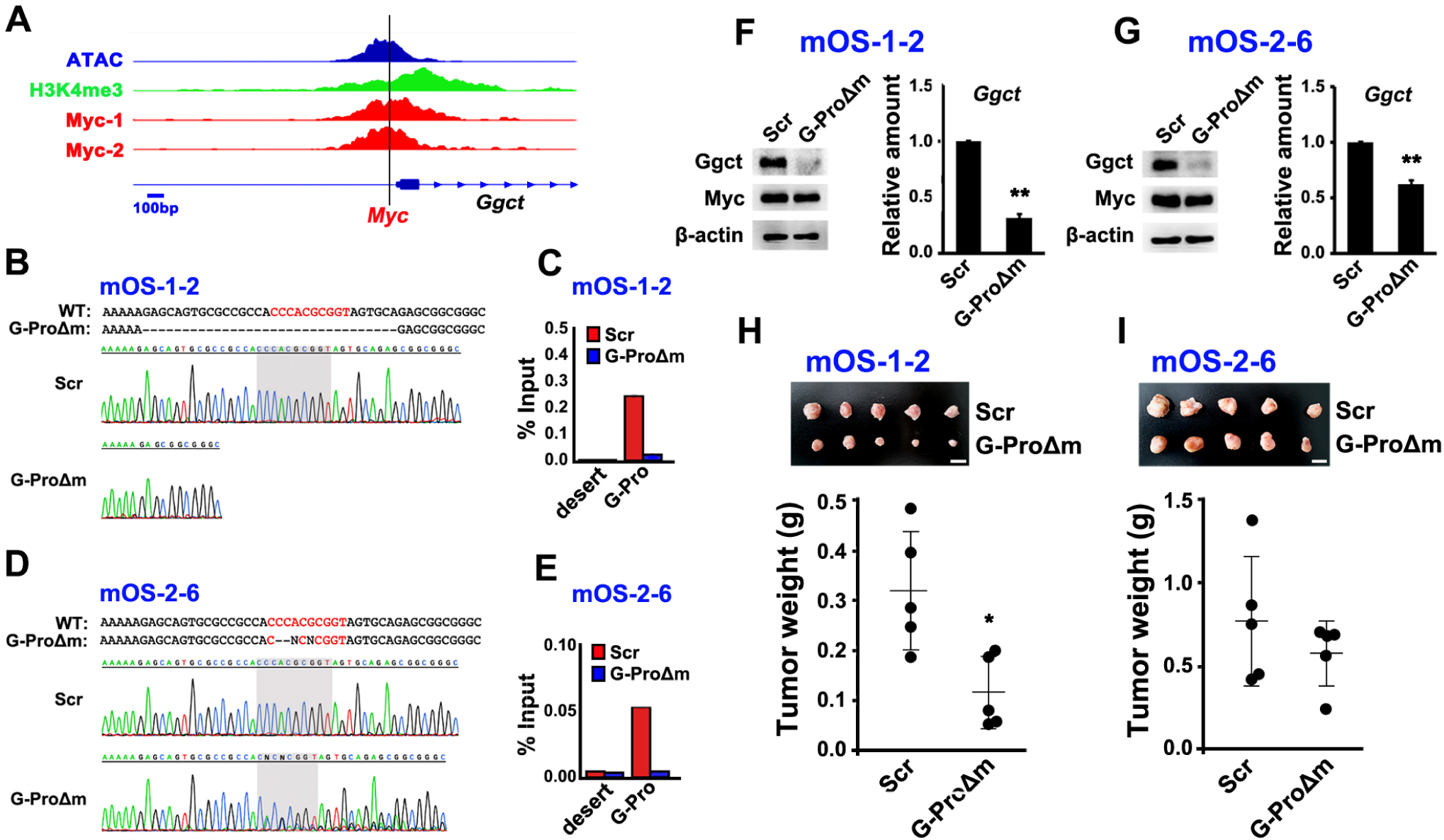
Upregulation of Ggct by Myc is critical for tumorigenicity of *p53*-deficient OS cells. (A) Profiles of open/active chromatin (ATAC and H3K4me3) in mouse OS cells (refs. SRX20246338 and SRX17122863, respectively) and Myc in mouse liver (Myc-1; ref. SRX1486521) and lung (Myc-2; ref. SRX7428137) tumor cells are aligned across the *Ggct* promoter region. The location of the Myc-consensus binding site is shown (*Myc*). (B) Sequence alignment of DNA from mOS-1-2 cells (the clonal cells shown as mOS-1 cl.2 in Figure 2D) from which the Myc-consensus site (shown in red) of the *Ggct* promoter was homologously deleted by genome editing (G-ProΔm) and non-targeted control mOS-1-2 cells (Scr). (C) ChIP-qPCR reveals occupancy of the Myc-consensus site of the *Ggct* promoter (G-Pro) in G-ProΔm and Scr mOS-1-2 cells by Myc. The gene desert region (desert) was amplified as a negative control. (D) Sequence alignment of DNA from mOS-2-6 cells (the clonal cells shown as mOS-2 cl.6 in Figure 2E) from which the Myc-consensus site (shown in red) of the *Ggct* promoter was homologously deleted by genome editing (G-ProΔm) and non-targeted control mOS-2-6 cells (Scr). (E) ChIP-qPCR reveals occupancy of the Myc-consensus site of the *Ggct* promoter (G-Pro) in G-ProΔm and Scr mOS-2-6 cells by Myc. The gene desert region (desert) was amplified as a negative control. (F and G) Amounts of the indicated proteins, as revealed by western blotting (left) and the relative amounts of *Ggct* mRNA, as revealed by qRT-PCR (right) in Scr and G-ProΔm of mOS-1-2 (F) and mOS-2-6 (G) cells. Data are presented as the mean ±SE (*n*=3). \*\**p*<0.01. (H and I) Tumorigenicity of Scr and G-ProΔm of mOS-1-2 (H) and mOS-2-6 (I) cells, as evaluated by allograft using nude mice (*n*=5). \**p*<0.05. Photos show tumors formed (H and I). Scale bars = 1 cm.

We used genome editing to delete the Myc-consensus site from the *Ggct* promoter (G-ProΔm) in clone 2 of mOS-1 (mOS-1-2) (Figure 5B) and in clone 6 of mOS-2 (mOS-2-6) (Figure 5D), both of which show marked tumorigenicity and expression of Myc and Ggct (Figures 2D, 2E, 4A and 4B). We found that Myc occupancy of the *Ggct* promoter in the G-ProΔm subclones of mOS-1-2 and mOS-2-6 cells fell to basal levels (Figure 5C and 5E). Correspondingly, expression of Ggct was reduced (Figure 5F and 5G) and their tumorigenicity was suppressed (Figure 5H and 5I) compared with controls (Scr).

Taken together, the present results indicate that the oncogenic potential of GGCT/Ggct in human and mouse osteosarcomagenesis. Binding of Myc to the *Ggct* promoter upregulated expression of Ggct, after which the latter plays a major role in elevating GSH levels in OS cells.

## DISCUSSION

In this study, we demonstrate that Ggct, an enzyme component of the γ-glutamyl cycle, is markedly upregulated in *p53*-deficient OS cells. We further show that Ggct is essential for the elevated levels of GSH observed and identify it as a novel direct target of Myc. Increased GSH synthesis, coupled with the resulting reduction in reactive oxygen species (ROS), is thought to confer a growth advantage on cancer cells^24^; indeed, GSH levels are elevated relative to normal tissues in a variety of human cancer types^12^.

The relationship between Myc activity and GSH levels has been studied primarily in cancer cells. The data show that GCL (GCLC and GCLM) are upregulated or downregulated, respectively, by MYC to increase or decrease GSH. Upon activation by ERK-dependent phosphorylation, MYC upregulates *GCLC* and *GCLM*^25^, also known as γ-glutamylcysteine synthetase (GCS) heavy and light subunits, respectively^11^. By contrast, *GCLC* is downregulated by a MYC-induced microRNA, miR-18a, in liver cancer cells^26^. Our analysis of a human OS cohort revealed a positive correlation between *MYC* and *GCLC* or *GCLM*, but this correlation was not as strong as that between *MYC* and *GGCT* (Figures 4E and S3). GGCT, recently identified as an enzyme belonging to the γ-glutamyl cycle, is becoming the target of in-depth investigations of its interaction with MYC, as well as the regulatory mechanisms, during cancer development.

Historically, Ras and Myc have been a paradigm for cooperation between oncogenes^27^. A previous study identified a crucial effector role of Myc; the paper shows that heterozygous deletion of Myc from a *KRas*^G12D^ and *p53*^R172H^-driven mouse model of pancreatic cancer markedly extends survival by confining development to the benign precursor stage^28^.Furthermore, introduction of a dominant-negative variant of Myc, termed Omomyc, leads to the regression of *KRas*^G12D^-driven lung adenocarcinomas^29^. Ras signaling has been shown to induce the phosphorylation of Myc at serine 62 stabilizing Myc and promoting its transcriptional activities^30^. The initial study that generated *Ggct-*knockout mice revealed oncogenicity of Ggct in the *KRas*^G12D^-driven lung cancer models^19^; this study clarified that oncogenic signals from Ras elevates Ggct expression markedly, which in turn, augments GSH levels. Here, we show that Myc, a transcription factor which was suggested to associate with oncogenic Ras in human OS cases^31^, is responsible for the upregulation of Ggct, and provide a comprehensive understanding of the mechanism of Ggct upregulation associated with tumorigenesis.

Previously, we described the oncogenic role of Runx3-Myc or Runx1-Myc axes in development of *p53*-defiecient OS^7–9^ or lymphoma^32^, respectively. In both cases, oncogenic Runx transcription factors upregulate Myc in the *p53*-deficient context^33^; thus it is possible that Runx3 upregulates Ggct. In fact, we found a significant correlation between expression of *RUNX3* and *GGCT* in the human OS cohort (R=0.25; p=0.022), but it was clearly weaker than the correlation between GGCT and MYC (Figure 4E). Furthermore, we found no significant occupation of the genomic region encoding the *GGCT/Ggct* gene by RUNX3/Runx3 in either humans or mice (data not shown). Therefore, we consider that Runx3 plays an indirect role in upregulation of Ggct, i.e., it acts via Myc.

Interestingly, knockdown of GGCT in several human cancer cell lines indirectly activates RB, leading to dephosphorylation of RB and cell growth arrest; this occurs through inactivation of the MEK-ERK pathway via downregulation of c-Met^34^. The tumor suppressor *RB* is the second most frequently mutated gene after *TP53* in sporadic human OS; indeed, gene alterations of *RB* were found in 31 of 86 patients in the TARGET cohort^7^. Inactivation of Rb facilitates development of *p53*-deficient OS in mice ^5,6,35^; thus if Ggct impairs the function of Rb indirectly, it could exert a marked osteosarcomagenesis-promoting function distinct from its role in increasing GSH levels.

To summarize, identified Ggct as a novel target of Myc during development of *p53*-deficient OS. The data suggest that Myc, via Ggct, maintains high GSH levels to promote osteosarcomagenesis. Since at least part of the function of oncogenic Myc is carried out by Ggct, targeting Ggct could offer a significant anti-tumor therapeutic strategy, either as an alternative to (or in conjunction with) targeting Myc, which is difficult to inhibit due to its wide range of functions.

### Limitations of the study

The study has several limitations. First, it is extremely challenging to examine the extent to which GSH itself is involved in the pathogenesis of OS. Furthermore, although we show that Ggct is crucial for elevation of GSH levels associated with the onset of OS, we did not examine other possible oncogenic functions of Ggct and its physiological functions in bone formation. In addition, since the present study used a conventional Cre mouse line, *Osx*-Cre, which does not allow for accurate cell lineage tracing^36^, the function of Ggct in the genuine cells of origin of OS is still unknown.

## MATERIALS AND METHODS

### Mouse lines

To generate the *Ggct*-targeted mouse line, a targeting vector harboring Loxp-flanked exon 2 and a FRT-flanked Neomycin resistance gene (Neo) was electroporated into Bruce-4 ES cells (C57BL/6). Digested genomic DNA isolated from targeted ES cells was subjected to Southern blotting using 5’ or 3’ alkaline phosphatase-labeled probes (490 bp or 422 bp), which were generated by PCR using primers; 5’- GTAGACAGGCCAATCCTGGTAGTTA -3’ and 5’-CAGCTAGTTCTCGACTAAGAAGCAC-3’, or 5’- GCTGAGTAGAACTGATAGCCTGGTA-3’ and 5’- GCAAACGTTAAAAGCACTTCATACT-3’, respectively, and mouse genome DNA as the template. To remove the FRT-flanked Neo, the offspring (F1) was crossed with *CAG*- FLP transgenic mice, and then was crossed again with *CAG*-Cre transgenic mice to generate *Ggct* heterozygous (*Ggct*^+/-^) mice. The *Ggct*-deleted and wild-type alleles of *Ggct*^+/-^ mice were detected by PCR using primers; 5’- TTGATCAGATCTCCTGATACTGGAA-3’ and 5’- CTTAGCATTTTGATATTGCAGTTGG-3’, which yielded products of 2031 bp and 204 bp, respectively.

A floxed *p53* mouse line was described previously^37^. The *Sp7/Osx*-Cre (no.006361) line was purchased from Jackson Laboratory. All mouse studies were performed in the C57BL/6 background using approximately equal numbers of males and females. The details of all animal experiments, including the number of mice (sample size) used, were reviewed and approved by the Animal Care and Use Committee of Nagasaki University Graduate School of Biomedical Sciences (no. 2104011709-4). Four mice were housed in each cage. Mice were reared in a pathogen-free environment under a 12 h light/dark cycle and a temperature of 22 ± 2°C.

### MSCs and mOS and human OS cells

All cells used in this study were confirmed to be free of mycoplasma infection and maintained in F12/DMEM supplemented with 10% fetal bovine serum. For the generation of BM-MSCs, BM cells were flushed from the femur of mice with F12/DMEM. Cd11b- and Cd45-negative adherent BM cells, which were negatively selected using a magnetic cell sorting system (MACS; Miltenyi Biotec) comprising of CD11b (no. 130-049-601) and CD45 (no. 130-052-301) MicroBeads and MS Columns (no. 130-042-201), were used as MSCs. Similarly, to generate mOS cells, adherent cells obtained from mouse OS tissues that had been minced and digested with collagenase I were negatively selected by MACS. Cd11b- and Cd45-negative OS cells were used as mOS cells. MNNG-HOS/Saos-2 and NOS1 cell lines were purchased from ATCC and RIKEN, respectively.

### Genome editing for deletion of a Myc-consensus binding site

The Myc-consensus binding site in the *Ggct* promoter was deleted by CRISPR-based genome editing. Briefly, HEK293T cells were co-transfected with a lentiCRISPRv2 plasmid (Addgene #52961) containing each sgRNA sequence (see below), the second-generation packaging plasmid psPAX2 (Addgene #12260), and the envelope plasmid pMD2.G (Addgene #12259), using the X-tremeGENE HP DNA Transfection Reagent (Roche). After filtration through a 0.45 μm filter, conditioned medium containing each lentivirus was used to infect mOS cells; infected cells were then selected with puromycin. After cloning resistant cells, cells harboring the intended deletion were identified by sequencing. The following sgRNA sequences were used: 5′- TGGTTTACATGTCGACTAAC-3′ (Scrambled; Scr) and 5′- GCAGTGCGCCGCCACCCACG-3′ (deletion of a Myc-consensus binding site in the *Ggct* promoter; G-ProΔm).

### shRNA-knockdown

Knockdown of endogenous gene expression was performed using the RNAi-Ready pSIREN-RetroQ Vector (Clontech). Briefly, cells were retrovirally transfected with shRNA and then selected with puromycin. Resistant cells were used for subsequent assays without cloning. The following shRNA sequences were used; Myc-1: 5′-GAACATCATCATCCAGGAC-3′ Myc-2: 5′-ACATCATCATCCAGGACTG-3′ MYC: 5′-AACAGAAATGTCCTGAGCAAT-3′ Luciferase (Luc) as a control: 5’-GTGCGTTGCTAGTACCAAC-3′

### RNA-seq

RNA-seq data and corresponding clinical information were obtained for 86 primary OS patients (hOS) from the TARGET project (https://ocg.cancer.gov/programs/target; accession number phs000468, NCBI dbGaP).) Of those, sequencing and clinical information were available for 84; these data were used for survival analysis. RNA-seq datasets from two human osteoblast (hOB) samples were derived from the ENCODE project under accession number GSE78608. RNA counts were quantified using Kallisto^38^ followed by pseudo-aligning FASTQ reads to the human genome (hg38). Survival analysis was performed after patients were stratified into low- and high-expressing groups based on the median value for each gene. Prior to correlation and survival analyses, human gene counts were normalized against the housekeeping gene *ACTB*, which encodes β-Actin. Correlation analysis (*GGCT* vs. *MYC*) was performed using Spearman’s correlation method. RNA-seq data from a previous study^9^, which were generated from MSCs and mOS cells isolated from three individual *OS* mice and submitted to DDJB sequence read archive with the accession number DRA012931, were used. Differential expression analysis of hOB versus hOS and MSCs versus mOS cells, was performed using edgeR, and the results were used to draw heatmaps.

### Immunoblotting and immunohistochemistry

Lysates of OS cells and MSCs were prepared for immunoblotting using a lysis buffer containing 9 M Urea, 2% Triton X-100, 2-mercaptoethanol, and protease/phosphatase inhibitors. Immunoblotting was performed using the following antibodies: anti-GGCT (16257-1-AP; proteintech), anti-c-Myc (ab32072; Abcam), anti-p53 (1C12; Cell Signaling Technology), and anti-β-actin (AC-15; Sigma-Aldrich).

OS tissues were fixed in 4% paraformaldehyde/PBS, decalcified in Decalcifying Soln. B (041-22031; Wako) at 4 ℃ for 5 days, embedded in paraffin, and cut into 4 µm sections. Anti-c-Myc (sc-764; Santa Cruz Biotechnology) and anti-GGCT (16257-1-AP; proteintech) antibodies were used for immunodetection on rehydrated sections pretreated with Target Retrieval Solution (S1699; DAKO). The Envision™+ system (HRP/DAB) (K4011; DAKO) was used for visualization. Counter staining was done with Mayer’s Hematoxylin Solution (131-09665; Wako).

### ChIP-qPCR, ATAC-seq and ChIP-Seq

ChIP experiments were performed using the SimpleChIP Enzymatic Chromatin IP kit and magnetic beads (Cell Signaling Technology). Briefly, 8 million cells were cross-linked for 10 min at room temperature with 1% formaldehyde. After permeabilization, cross-linked cells were digested with micrococcal nuclease and then immunoprecipitated with an anti-Myc antibody (ab32072; Abcam) or normal rabbit IgG (#2729; Cell Signaling Technology). Immunoprecipitated products were isolated with Protein G Magnetic Beads (Cell Signaling Technology) and subjected to reverse cross-linking. DNA was subjected to quantitative PCR (qPCR) using primer pairs targeting the following; mouse gene desert: 5′-ACCAAGCACAGAAAAGGTTCAAAC-3′ and 5′-TCCAGATGCTGAGAGAAAAACAAC-3′; human gene desert: 5’- TGAGCATTCCAGTGATTTATTG-3’ and 5’-AAGCAGGTAAAGGTCCATATTTC-3’; Myc-consensus binding site in the mouse *Ggct* promoter: 5′- ACAGAGAAGCCGGACTAGCG-3′ and 5′-AAGCCGGCAGCCAATCCTC-3′; MYC-consensus binding site in the human *GGCT* promoter: 5′- GCCAGAGAGCGCAACACTGG-3′ and 5′-AGCGCTCGCTCCTGACTCG-3′; and MYC-consensus binding site in the human *CDK4* promoter: 5′- ACACCTCTGCTCCTCAGAGC-3′ and 5′-AGGAGGGCGAAGAGTGTAAGG-3′. Real-time quantitative PCR reactions were performed on a 7300 Real-time PCR system (ABI) using THUNDERBIRD SYBR qPCR Mix (Toyobo).

ATAC-seq and ChIP-seq data were obtained from ChIP-Atlas (https://chip-atlas.org) and analyzed.

### qRT-PCR

Total RNA was extracted using the NucleoSpin RNA kit (Macherey-Nagel) and reverse-transcribed into cDNA using the ReverTra Ace qPCR RT Master Mix (Toyobo). Real-time quantitative PCR reactions were performed on a 7300 Real-time PCR system (Applied Biosystems) using the THUNDERBIRD SYBR qPCR Mix (Toyobo) and the following primer sets; *Ggct*: 5′-GTACTTCGCCTACGGCAGCA-3′ and 5′-CTTCGTCGCCAGGACTTTGA-3′ and *Actb*: 5′-CATCCGTAAAGACCTCTATGCCAAC-3′ and 5′- ATGGAGCCACCGATCCACA-3′

### Tumorigenicity of cells

Transplantation (allografts) was performed by subcutaneous injection of mOS cells (5 × 10^6^) cells into 6–8-week-old female BALB/c-*nu/nu* mice (nude mice). Tumorigenicity was assessed by measuring tumor weight at 45 days post-inoculation.

### Micro (μ)-CT analysis

µCT analysis was performed using a µCT system (R_mCT; Rigaku Corporation, Tokyo). Data from scanned slices were used for three-dimensional analysis to calculate femoral morphometric parameters. Trabecular bone parameters were measured at the distal femoral metaphysis. Craniocaudal scans of approximately 2.4 mm (0.5 mm from the growth plate) were obtained for 200 slices in 12 μm increments. Cortical bone parameters were measured at the mid-diaphysis of the femur. The threshold mineral density was 500 mg/cm^3^.

### Measurement of GSH

GSH levels was measured using a GSSG/GSH Quantification kit (G257; DOJINDO Laboratories). Briefly, cells were frozen and thawed in 10 mM HCl and then mixed with 5% 5-sulfosalicylic acid dihydrate (190–04572; Wako). After centrifugation, GSH/GSSG in the supernatant were detected by measuring the absorption derived from the colorimetric reaction between DTNB and the enzymatic recycling system. The concentration of GSH and GSSG was read from a standard glutathione calibration curve and the GSH concentration was determined by subtracting the concentration of GSSG from total GSSG/GSH concentration. Luminescence was measured at 405 nm in a Multiskan™ GO spectrophotometer (Thermo Fisher Scientific), and the values were normalized to cell numbers.

### Statistics

All quantitative data are expressed as the mean ± SEM. Differences between groups were analyzed using an unpaired two-tailed Student’s t-test (two groups) or one-way analysis of variance (more than two groups). All analyses were performed using Prism 8 (GraphPad software). Survival was analyzed using the Kaplan–Meier method and data compared using the log-rank test. A p value of <0.05 was deemed significant. No samples from *in vivo* and *in vitro* experiments were excluded from the analysis.

## ACKNOWLEDGMENTS

We thank A. Berns for providing the p53 flox mouse line, S. Tanaka for technical help, and all members of Research Center for Biomedical Models and Animal Welfare, Nagasaki University for maintaining the mouse lines. We are deeply grateful to Prof Tatsuhiro Yoshiki, whose early contributions to this project were invaluable. Though no longer with us, his influence endures in our work.

This work was supported by KAKENHI/Japan Society for the Promotion of Science (JSPS) grants 18H02972 (K.I.), 19K22724 (K.I.), and 21H03113 (K.I.), and by the Funding Program for Next Generation World-Leading Researchers LS097 (K.I.).

## AUTHOR CONTRIBUTIONS

K.I. initiated the study. T.U., S.O., Y.D., and K.I. designed the experiments. T.U, S.O., Y.D, Y.K., Y.N., T.I., and K.I. conducted the experiments. T.U and Y.D. performed bioinformatic analyses. T.U., S.O., and K.I. wrote the manuscript. H.I., S.K., and S.N. provided experimental information and coordinated the project. K.I. supervised the study.

