## Supplementary information for "Myc upregulates Ggct, γ-glutamylcyclotransferase to promote development of *p53*-deficient osteosarcoma"

Running title: Myc upregulates Ggct in *p53*-deficient OS

**This file includes:**

**Supplementary Figures 1-5**

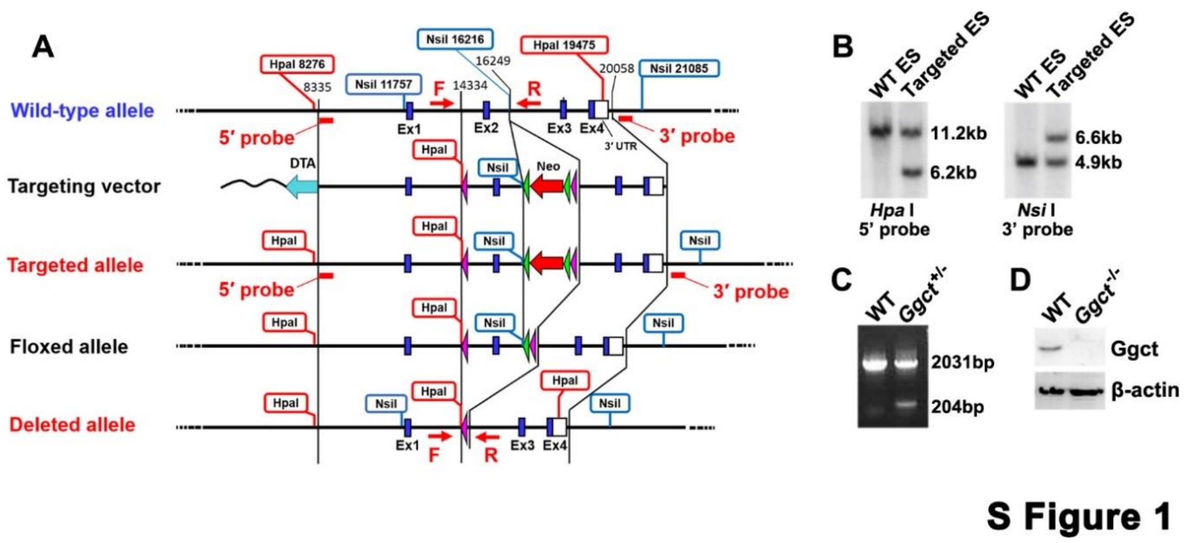


**Supplementary Figure 1. Generation of *Ggct*-deleted mice (related to Figure 3A)**

(A) Targeting vector used for homologous recombination, and the strategy used to generate the *Ggct*-deleted mouse line.

(B) Southern blot analysis. Genomic DNA from wild-type and targeted ES cells was digested with *Hpa* I (left) or *Nsi* I (right) and detected using the 5' or 3' probes shown in (A).

(C) Genotyping PCR using the F and R primers shown in (A) to detect wild-type (WT) and *Ggct*-deleted alleles.

(D) Western blotting shows that Ggct was not expressed in MSCs isolated from *Ggct*$^{-/-}$ mice.

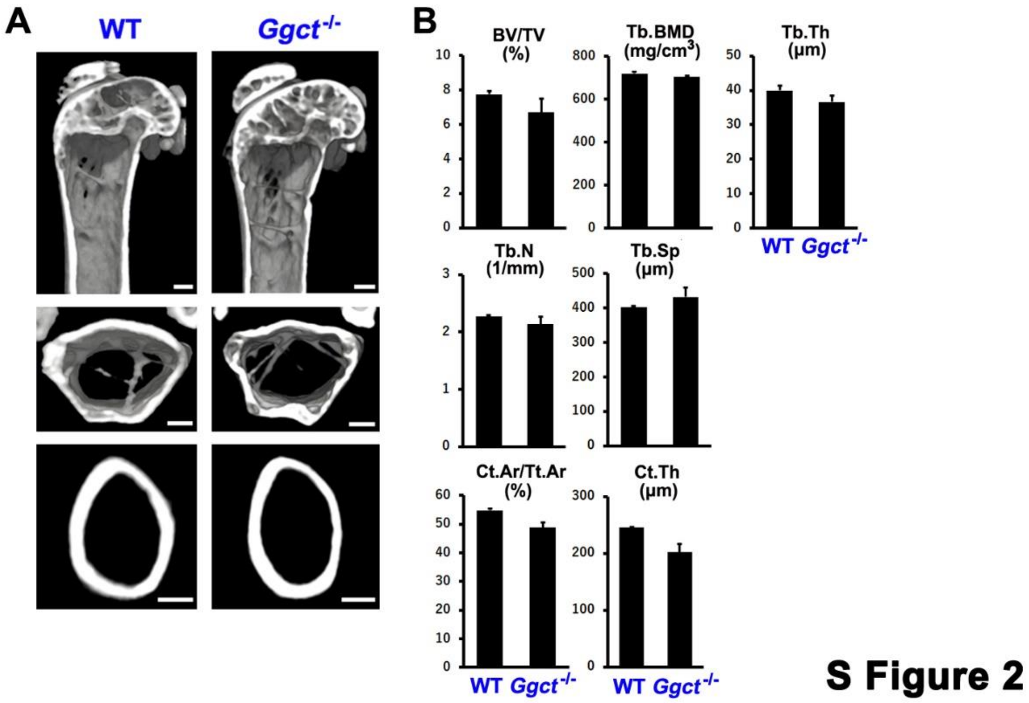


**Supplementary Figure 2. Bone formation in the femur of adult female wild-type and *Ggct*$^{-/-}$ mice (related to Figure 3D and 3E)**

(A) Representative μCT three- or two-dimensional images of the bone architecture of female wild-type (WT) and *Ggct*$^{-/-}$ mice aged 15 months. Images of trabecular bone in the distal femoral metaphysis (upper and middle), and cortical bone at mid-diaphysis, in the femur (lower) are shown. Scale bars = 500 μm.

(B) Quantification of the trabecular bone volume (bone volume/tissue volume, BV/TV), trabecular bone mineral density (Tb.BMD), trabecular thickness (Tb.Th), trabecular number (Tb.N), trabecular separation (Tb.Sp), cortical area (Ct.Ar/Tt.Ar), and cortical thickness (Ct.Th) in female WT and *Ggct*$^{-/-}$ mice aged 15 months. Data are presented as the mean ± SE (n=3).

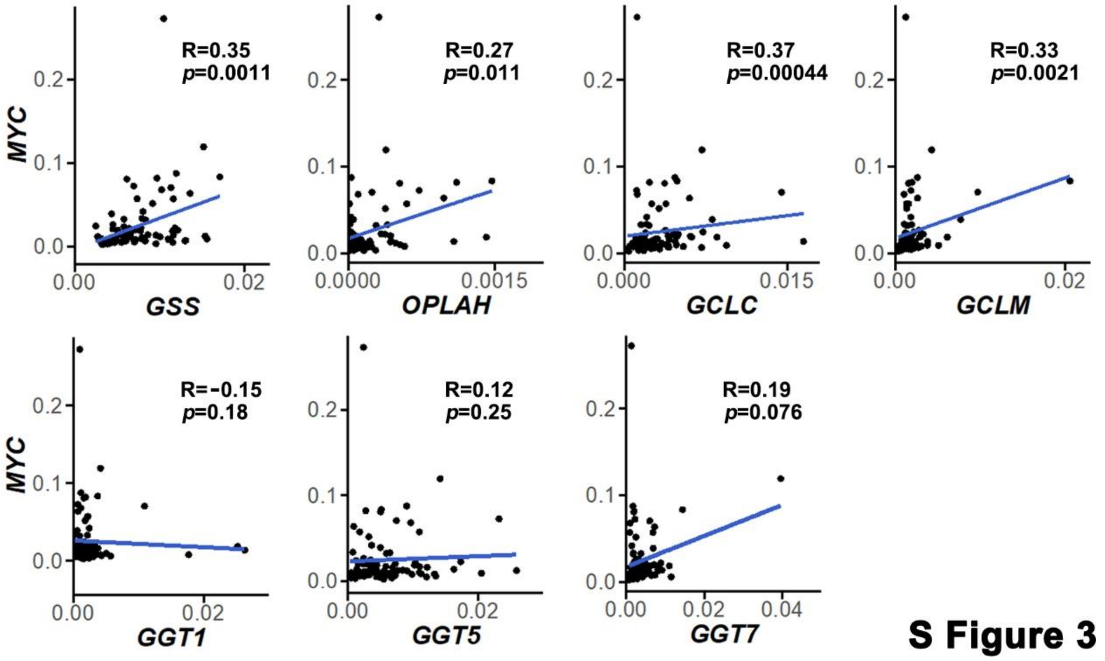


**Supplementary Figure 3. Correlation between expression of *MYC* and genes encoding enzymes in the γ-glutamyl cycle (related to Figure 4E).**

MYC expression was plotted against that of various genes encoding γ-glutamyl cycle enzymes in cases of human OS (n=86, Figure 1B). Spearman's rank correlation coefficient (R) is shown.

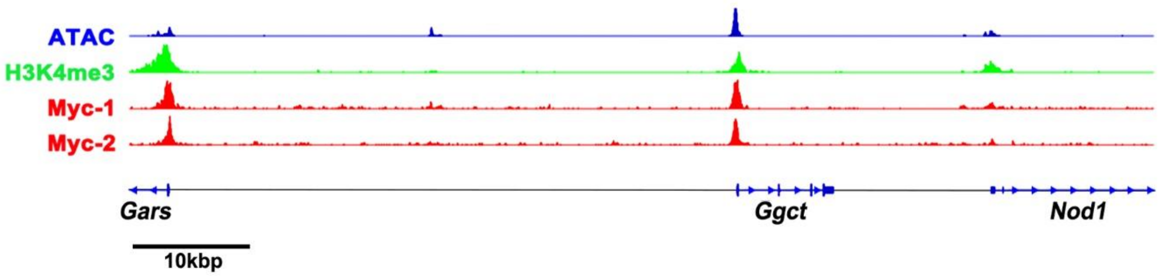


**S Figure 4**

**Supplementary Figure 4. Profiles of open/active chromatin and Myc in mouse genomic regions surrounding *Ggct* (related to Figure 5A)**

Profiles of open/active chromatin (ATAC and H3K4me3) in mouse OS cells (refs. SRX20246338 and SRX17122863, respectively) and Myc in mouse liver (Myc-1; ref. SRX1486521) and lung (Myc-2; ref. SRX7428137) tumor cells, are aligned across the region of the mouse genome (~100kbp) containing the entire *Ggct* gene.

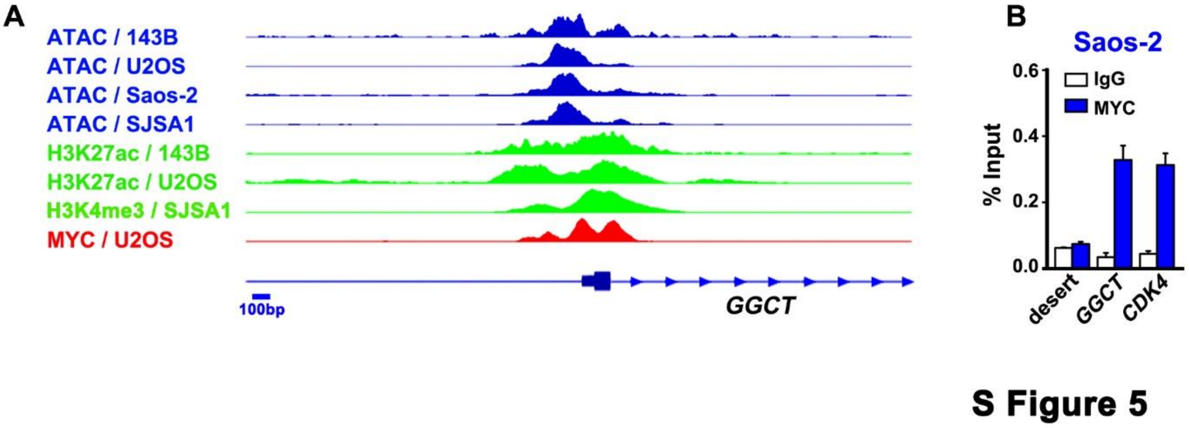


**Supplementary Figure 5. Profiles of open/active chromatin and MYC in the human *GGCT* promoter (related to Figure 5A)**

(A) Profiles of open/active chromatin (ATAC and H3K27ac/H3K4me3) and Myc in the following human OS cell lines are aligned within the *GGCT* promoter region; 143B (ATAC; SRX10169934 and H3K27ac; SRX5975304), U2OS (ATAC; SRX7030826, H3K27ac; SRX10829244 and MYC; SRX8299829), Saos-2 (ATAC; SRX7644740) and SJSA1 (ATAC; SRX7644737 and H3K4me3; SRX10187663).

(B) Occupancy of the human *GGCT* promoter by MYC, shown with a positive control, the promoter of *CDK4* which is a known target of MYC, and a negative control, gene desert (desert) in Saos-2 cells, as revealed by ChIP-qPCR. IgG was used as a negative control for ChIP. Data are presented as the mean ±SE ($n$=3).
